# c-di-AMP hydrolysis by a novel type of phosphodiesterase promotes differentiation of multicellular bacteria

**DOI:** 10.1101/789354

**Authors:** Andreas Latoscha, David Jan Drexler, Mahmoud M. Al-Bassam, Volkhard Kaever, Kim C. Findlay, Gregor Witte, Natalia Tschowri

## Abstract

Antibiotic-producing *Streptomyces* use the diadenylate cyclase DisA to synthesize the nucleotide second messenger c-di-AMP but the mechanism for terminating c-di-AMP signaling and the proteins that bind the molecule to effect signal transduction are unknown. Here, we identify the AtaC protein as a new type of c-di-AMP-specific phosphodiesterase that is also conserved in pathogens such as *Streptococcus pneumoniae* and *Mycobacterium tuberculosis*. AtaC is monomeric in solution and binds Mn_2+_ to specifically hydrolyze c-di-AMP to AMP via the intermediate 5’-pApA. As an effector of c-di-AMP signaling, we characterize the RCK-domain protein CpeA as the first c-di-AMP-binding protein to be identified in *Streptomyces*. CpeA interacts with the predicted cation / proton antiporter, CpeB, linking c-di-AMP signaling to ion homeostasis in actinobacteria. Hydrolysis of c-di-AMP is critical for normal growth and differentiation in *Streptomyces*, connecting osmotic stress to development. Thus, we present the discovery of two novel components of c-di-AMP signaling in bacteria and show that precise control of this second messenger is essential for osmoregulation and coordinated development in *Streptomyces*.

## INTRODUCTION

Bacteria use mono-, di-, and trinucleotides as second messengers to control fundamental physiological functions in response to signal sensing (1). Among these molecules, cyclic di-3’,5’-adenosine monophosphate (c-di-AMP) is the only nucleotide messenger that must be precisely balanced, since both, its depletion and overproduction can be toxic (2). Its core function is to control cellular integrity by setting homeostasis of osmolytes that in many bacteria are used for osmoregulation (3, 4). Changes of external osmolarity trigger water fluxes across the membrane, which can lead to cell dehydration or swelling and finally collapse or burst when osmobalance mechanisms fail to respond properly (5). As a key component of these mechanisms, c-di-AMP directly targets transport systems for osmoactive and osmoprotective substances such as potassium ions and low-molecular-weight compatible solutes in many bacteria (6-10).

c-di-AMP also plays a central role in host-pathogen interactions and bacterial virulence (11). Secreted c-di-AMP is recognized by host’ innate immunity receptors STING, DDX41 and RECON to regulate type I interferon immune response and NF-kB pathways, respectively (12-15). Modulation of intracellular c-di-AMP has been reported to affect virulence of *Streptococcus pyogenes* (16), *Listeria monocytogenes* (17), *Streptococcus pneumonia* (18) and *Mycobacterium tuberculosis* so that the molecule is considered as an attractive antimicrobial target (19).

c-di-AMP synthesis out of two ATP molecules is catalyzed by the diadenylate cyclase (DAC) activity of the DisA_N domain (Pfam PF02457), which was identified in the structural and biochemical analysis of the DNA-integrity scanning protein A (DisA) of *Thermotoga maritima* (20). DisA is mainly present in sporulating firmicutes and actinobacteria (21) and has a conserved domain organization consisting of a N-terminal DAC domain and a C-terminal DNA-binding helix-hairpin-helix domain separated by a linker region (20). C-di-AMP hydrolysis is mediated by the DHH-DHHA1 domain containing the Asp-His-His motif. The multidomain membrane-associated GdpP protein in *Bacillus subtilis* was the first characterized DHH-DHHA1-type phosphodiesterase (PDE) (22). In addition, HD domains with a catalytic His-Asp motif, which were first identified in the PgpH protein in *L. monocytogenes*, also degrade c-di-AMP (17).

However, most actinobacteria contain DisA for c-di-AMP synthesis but do not encode DHH-DHHA1-domain containing or HD-type c-di-AMP PDEs. Hence, we wondered how actinomycetes balance intracellular c-di-AMP. Within actinobacteria, *Streptomyces* are the most extensively studied mycelial organisms and the richest natural source of antibiotics (23). For growth and reproduction, *Streptomyces* undergo a complex developmental life cycle, which involves the conversion between three morphologically and physiologically distinct forms of cell existence. During exponential growth, they proliferate by extension and branching of vegetative hyphae. The switch to stationary phase and onset of the reproductive phase is marked by the erection of aerial hyphae. These filaments elongate and divide into unigenomic prespore compartments that ultimately mature into chains of spores. Completion of the developmental program is easily visible by eye since mature *Streptomyces* spores accumulate a spore pigment. For example, our model species, the chloramphenicol producer *S. venezuelae*, is characterized by a green spore pigment such that colonies turn green at the end of the life cycle (24, 25). Importantly, antibiotic production and morphological differentiation are co-regulated in *Streptomyces*. Hence, studying their developmental biology also provides a better understanding of the control of their secondary metabolism.

In this work, we identified and characterized the PDE superfamily protein AtaC as the founding member of a novel type of c-di-AMP-specific hydrolases. AtaC is broadly distributed in bacteria and the only known c-di-AMP PDE in most actinomycetes. Among others, pathogens such as the causative agent of pneumonia, *S. pneumoniae*, contain an AtaC homolog that we characterize here to be a functional c-di-AMP hydrolase. Our biochemical and structural analyses show that AtaC is a monomeric Mn_2+_-dependent PDE with high affinity for c-di-AMP. Moreover, we provide direct biochemical evidence that *Streptomyces* DisA is an active DAC and that c-di-AMP produced by DisA is crucial for survival under ionic stress conditions. Further, we show that accumulation of c-di-AMP in the *S. venezuelae ataC* mutant results in profound developmental and growth defects and report the identification of the RCK_C-domain (RCK for regulator of conductance of K_+_) containing protein CpeA as the first c-di-AMP binding protein in *Streptomyces*. Overall, in this study we identified and functionally characterized core components of c-di-AMP signaling in *Streptomyces* and link c-di-AMP regulation with ion homeostasis to control differentiation in multicellular bacteria.

## RESULTS

### DisA is the major c-di-AMP synthetase in *S. venezuelae*

DisA is the sole DAC protein encoded in the *S. venezuelae* genome and is conserved in all sequenced *Streptomyces* strains. To demonstrate DisA DAC activity, we purified N-terminally his-tagged DisA and DisA_D86A_ that carries an alanine instead of aspartate in the active site. We included his-tagged *B. subtilis* DisA (DisA_Bsu_) as a positive control for enzymatic activity (20). [_32_P]-labeled ATP was added as substrate for *in vitro* DAC assays and the reactions were separated by thin layer chromatography (TLC). DisA synthesized c-di-AMP whereas the mutated DisA_D86A_ failed, demonstrating that *S. venezuelae* DisA is a functional DAC, which requires the conserved catalytic aspartate D_86_ for activity (Figure 1A).

**Figure 1.**
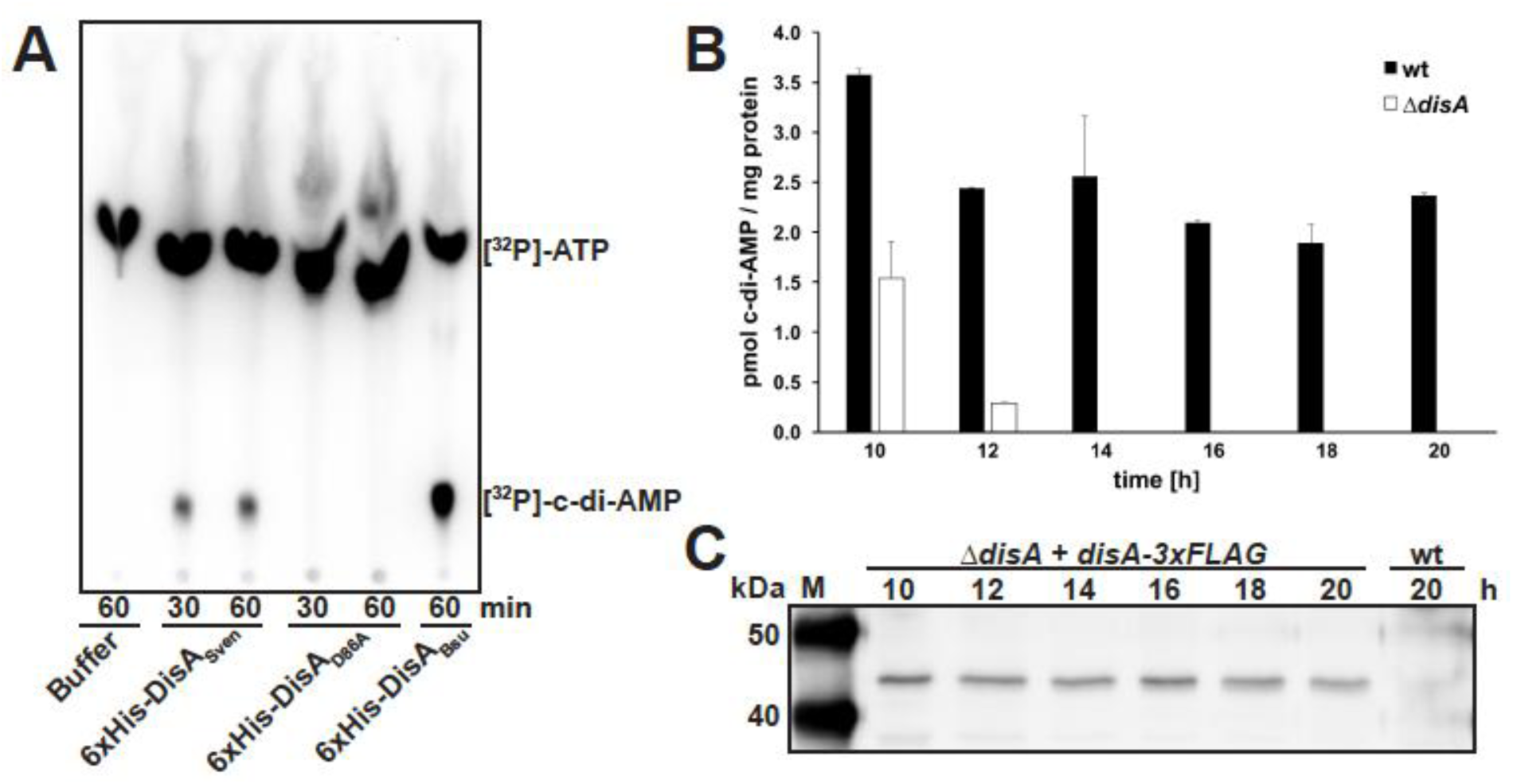
DisA is an active diadenylate cyclase *in vitro* and *in vivo*. (A) Thin layer chromatography of diadenylate cyclase (DAC) assay with purified 6xHis-DisA_Sven_ and 6xHis-DisA_D86A_, and [_32_P]-ATP as substrate. Migration of [_32_P]-ATP in buffer is shown in lane 1. 6xHis-DisA_Bsu_ served as positive control for DAC activity. (B) Intracellular c-di-AMP levels in *S. venezuelae* wild type and *ΔdisA* during late vegetative growth (10 to 12 h), early sporulation (14 to 16 h) and sporulation (from 18 h). Data are presented as mean of biological replicates ± standard deviation (n=3). (C) Expression profile of DisA-3xFLAG in a *disA* mutant complemented with *disA-3xFLAG* under control of *disA* promoter grown in liquid sporulation medium (MYM). DisA-3xFLAG was detected using a monoclonal anti-FLAG antibody. Wild type served as negative control.

*In vivo*, DisA is the major source for c-di-AMP in *S. venezuelae* (Figure 1B) (26), however, we reproducibly detected low c-di-AMP levels in Δ*disA* during vegetative growth (10 and 12 h), which disappeared upon onset of sporulation (14 h), suggesting that *S. venezuelae* might contain a non-DAC-domain enzyme capable of c-di-AMP production (Figure 1B). The presence of c-di-AMP throughout the wild type *S. venezuelae* life cycle suggested that *disA* expression is constitutive. To confirm this, we complemented the *disA* mutant by chromosomal insertion of a C-terminally 3xFLAG-tagged *disA* under control of its native promoter. Using a monoclonal anti-FLAG antibody, we detected constant DisA-3xFLAG expression in all developmental stages, which correlated with c-di-AMP production in the wild type under the conditions tested (Figure 1C).

Altogether, our data show that DisA is a functional DAC *in vitro* and *in vivo* and the major enzyme for c-di-AMP production in *S. venezuelae*.

### The phosphodiesterase superfamily protein AtaC (Vnz_27310) degrades c-di-AMP

*Streptomycetes* do not possess PDEs with a DHH-DHHA1 domain or a PgpH-type HD domain, known to degrade c-di-AMP in other bacteria (17, 22), raising the question as to how *S. venezuelae* removes c-di-AMP from the cytoplasm. To find a potentially novel c-di-AMP PDE, we used interproscan (http://dx.doi.org/10.7717/peerj.167) to search for Pfam PF01663, which is associated with putative type I phosphodiesterases/nucleotide pyrophosphatases. Among others, we found two proteins (Vnz_27310 and Vnz_31010) belonging to the phosphodiesterase and metallophosphatase superfamilies, respectively, that we selected for *in vitro* PDE activity tests.

Purified N-terminally his-tagged Vnz_27310 and Vnz_31010 were assayed *in vitro* using [_32_P]-labeled c-di-AMP as substrate. While we could not detect [_32_P]-c-di-AMP cleavage activity for Vnz_31010, Vnz_27310 clearly degraded c-di-AMP to 5′-pApA and finally to AMP (Figure 2A) so that we named Vnz_27310 AtaC for <u>a</u>ctinobacterial PDE <u>ta</u>rgeting <u>c</u>-di-AMP. Addition of unlabeled c-di-AMP but not of c-di-GMP or cAMP competed with [_32_P]-c-di-AMP and led to reduced cleavage of the radiolabeled substrate, showing specificity for c-di-AMP (Figure 2A). We analyzed the kinetics of c-di-AMP hydrolysis activity of Vnz_27310 using anion exchange chromatography assays and determined a *k*_cat_ of 0.2 s_-1_ (Figure S1 A-B), while only a negligible c-di-GMP hydrolysis activity was detected (Figure S1 C). We also compared Vnz_27310-dependent hydrolysis of the linear dinucleotides 5′-pApG and 5′-pGpG to the hydrolysis of 5′-pApA and observed a high hydrolysis activity for 5′-pApA (k_cat_= 2.1 s_-1_), whereas the other substrates tested were only degraded to a small extent (Figures 2B and S1 D-F).

**Figure 2.**
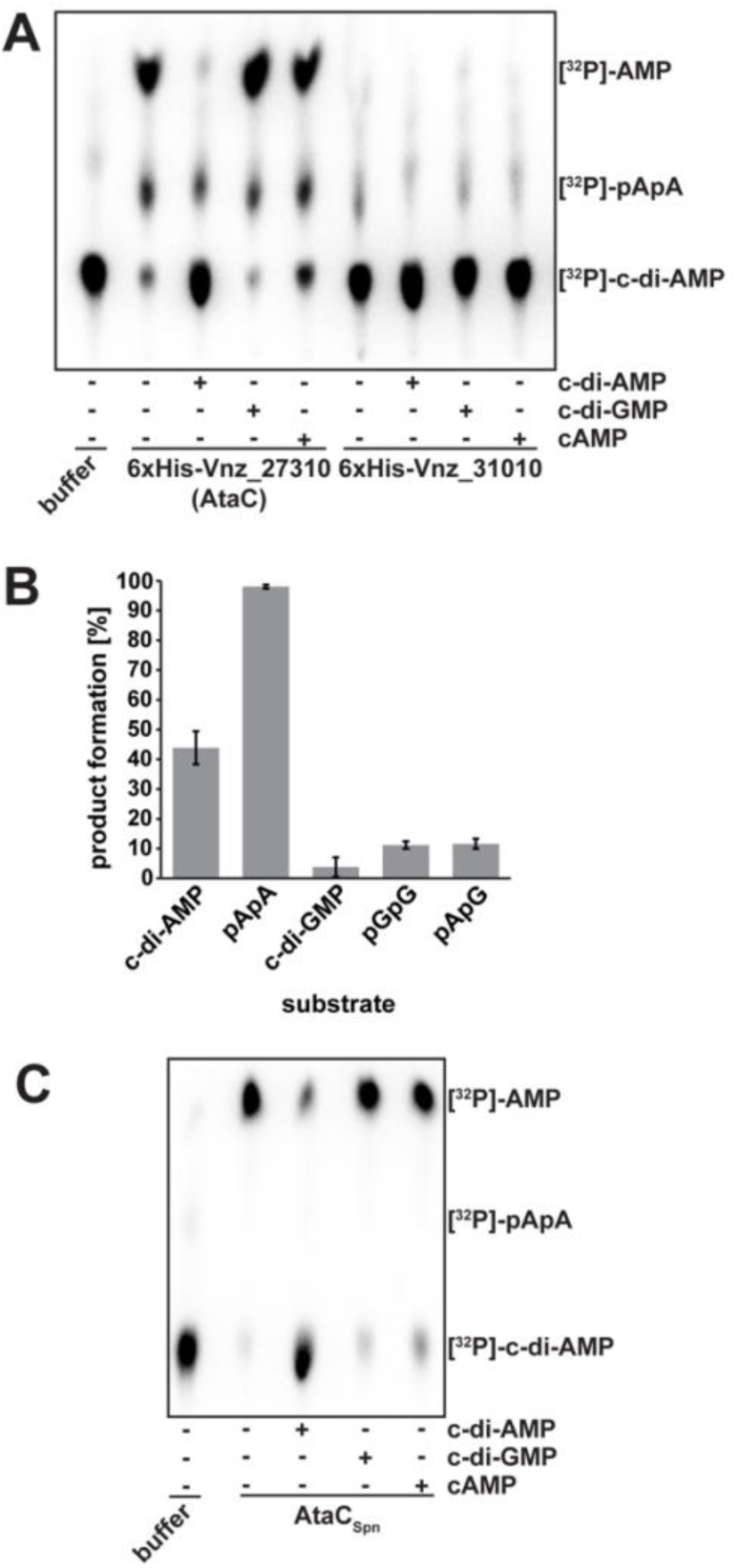
AtaC is a c-di-AMP-specific phosphodiesterase (PDE). Thin layer chromatography of PDE assay of AtaC and Vnz_31010 from *S. venezueale* (A) and *S. pneumoniae* (AtaC_SPN_) (C) with [_32_P]-c-di-AMP. Radioactively labeled c-di-AMP in buffer migrates as shown in lane 1. In samples used for competition, unlabeled c-di-AMP, c-di-GMP or cAMP (indicated by “+”) were added in excess before starting the reaction with [_32_P]-c-di-AMP. (B) AtaC activity assay by ion-exchange chromatography runs on a 1 ml Resource Q column of the reaction products after 1 h incubation from 100 µl reactions containing 100 nM AtaC + 250 µM c-di-AMP, 5′-pApA, c-di-GMP, 5′-pGpG or 5′-pApG (n=3).

Using the PATRIC database (https://www.patricbrc.org), we examined the distribution of the here discovered c-di-AMP PDE (PGF_00172869) and found at least 5374 prokaryotic species containing homologs to AtaC (Table S2), including pathogens such as *S. pneumoniae* and *M. tuberculosis*. AtaC from *S. pneumoniae* (AtaC_SPN_; sequence ID: CVN04004.1) and from *M. tuberculosis* (AtaC_MTU_; sequence ID: CNE38097.1) share 41 % and 47 %, respectively, identical residues with AtaC from *S. venezuelae*. In agreement with the high degree of protein identity, enzyme assays data shown in Figure 2C demonstrate that AtaC_SPN_ also represents a c-di-AMP PDE and AtaC_MTU_ likely has the same function.

In summary, we identified and functionally characterized the sole c-di-AMP hydrolase in *Streptomyces* and a new c-di-AMP signaling component in pathogens and show that AtaC is a conserved phosphodiesterase that efficiently and specifically hydrolyzes c-di-AMP to AMP via the intermediate 5′-pApA.

### AtaC is a monomeric Mn_2+_-dependent phosphodiesterase

To further characterize the c-di-AMP hydrolysis mechanism of AtaC and to gain some structural insights into this PDE, we used HHpred (27) and found two close structural homologs. The core domain of a phosphonoacetate hydrolase (PhnA) from *Sinorhizobium meliloti* 1021 (28) and PDB code 3SZY) showed highest similarity and served as a template for the structural model of AtaC including the putative active site. The predicted active site comprises three aspartates (D68, D227 and D269), three histidines (H231, H270 and H384) and one threonine (T108) (Figure 3A).

**Figure 3.**
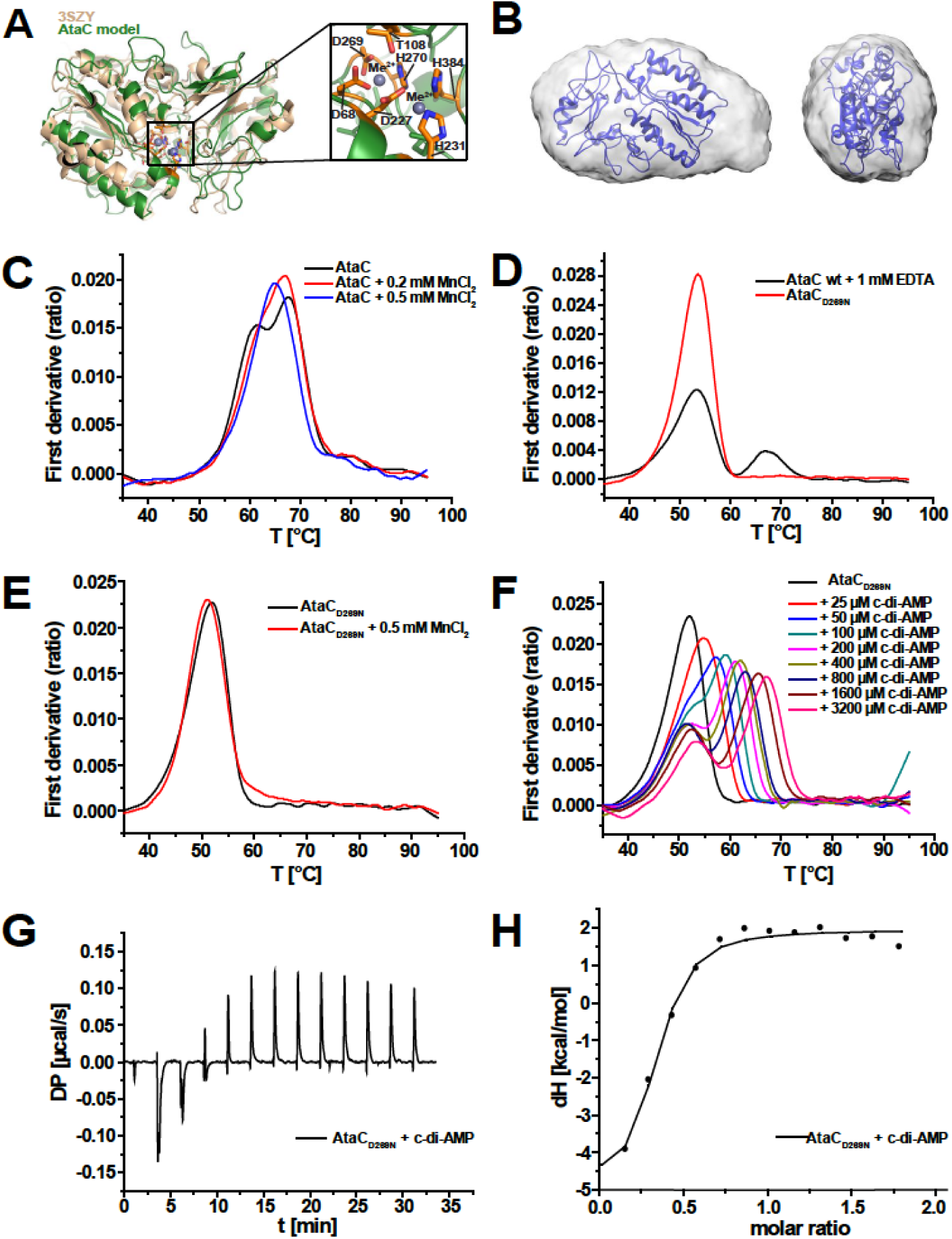
AtaC is a monomeric Mn_2+_-dependent phosphodiesterase. (A) Model of AtaC obtained from HHpred/MODELLER (green) superimposed with best match 3SZY (beige), Zoom-In shows the predicted active site, annotated with all most conserved residues. (B) Modelled structure from (A) superimposed with the final averaged and filtered *ab initio* shape (16 ab initio models averaged) from SEC-SAXS with front view (left) and side view (right). (C) nanoDSF thermal shift first derivative curves of 10µM apo AtaC (black), 10µM AtaC + 0.2 mM MnCl_2_ (red) and 10µM AtaC + 0.5 mM MnCl_2_ (blue). (D) nanoDSF thermal shift first derivative curves of 10 µM AtaC + 1 mM EDTA (black) and 10µM AtaC_D269N_ (red). (E) nanoDSF thermal shift first derivative curves of 10 µM AtaC_D269N_ (black) and AtaC_D269N_ + 0.5 mM MnCl_2_ (red). (F) nanoDSF thermal shift first derivative curves of 10 µM AtaC_D269N_ + c-di-AMP (25 µM - 3200 µM). (G) ITC measurement raw data of 23 µM AtaC_D269N_ mutant titrated with 231 µM c-di-AMP. (H) Binding curve and fit of ITC titration of the AtaC_D269N_ mutant with c-di-AMP (K_D_ = 731 ± 266 nM) (n=3).

Our size-exclusion chromatography (SEC) coupled multi-angle laser light scattering (MALLS) data show that AtaC is a monomer in solution with a molecular weight of 43.7 kDa (Figure S2A). The calculated *ab initio* shape of AtaC from SEC-SAXS (size-exclusion coupled small-angle X-ray scattering) data superimposes well with the HHpred model structure (Figure 3B) and the measured SAXS curve of AtaC is very similar to the theoretical scattering curve of PhnA (Figure S2 B-D), indicating that AtaC and PhnA have a similar shape in solution.

The enzymatic reaction of the PhnA-class hydrolases is known to be catalyzed by two metal ions in the active site (28) so we tested metal binding for AtaC by thermal unfolding assays using nano differential scanning fluorimetry (nanoDSF) assay and observed protein stabilization upon addition of manganese ions (Mn_2+_) (Figure 3C). Based on the structural similarity to PhnA, we identified potential metal-binding residues in AtaC and generated a variant, AtaC_D269N_, that we expected to lack Mn_2+_ coordination but retain nucleotide binding, as shown for DHH-DHHA1-type PDEs (22, 29). NanoDSF data confirmed stability of AtaC_D269N_ with a melting temperature comparable to the wild type protein when incubated with ethylenediaminetetraacetic acid (EDTA) (Figure 3D). Moreover, AtaC_D269N_ behaved identically to the wild type protein during purification and final SEC. In line with our predictions, AtaC_D269N_ failed to bind Mn_2+_ (Figure 3E) and did not hydrolyze c-di-AMP, as shown using ion exchange chromatography (IEX) based assays (Figure S3A). However, AtaC_D269N_ was still capable of c-di-AMP binding, as confirmed by nanoDSF experiments that showed a shift in the melting curve with increasing ligand concentration (Figure 3F). Using isothermal titration calorimetry (ITC) analysis we determined the dissociation constant (*K*_d_) of AtaC_D269N_ for c-di-AMP to be 731 ± 266 nM, whereas binding of c-di-GMP could not be detected (Figures 3G-H, and Figure S3B).

Altogether, our combined structural analysis and biochemical data strongly suggest that AtaC uses the same metal-ion dependent mechanism as its structural homolog PhnA for substrate cleavage.

### AtaC hydrolyzes c-di-AMP *in vivo*

We quantified c-di-AMP in cell extracts isolated from wild type *S. venezuelae* and the *ataC* null mutant using LC-MS/MS. Our data show that c-di-AMP levels are elevated in the *ataC* mutant during all developmental stages when compared to the wild type, demonstrating that AtaC degrades c-di-AMP *in vivo* and thus is an important component of c-di-AMP metabolism in *S. venezuelae* (Figure 4A). Western blot analysis showed that AtaC is constitutively expressed across the developmental cycle (Figure 4B).

**Figure 4.**
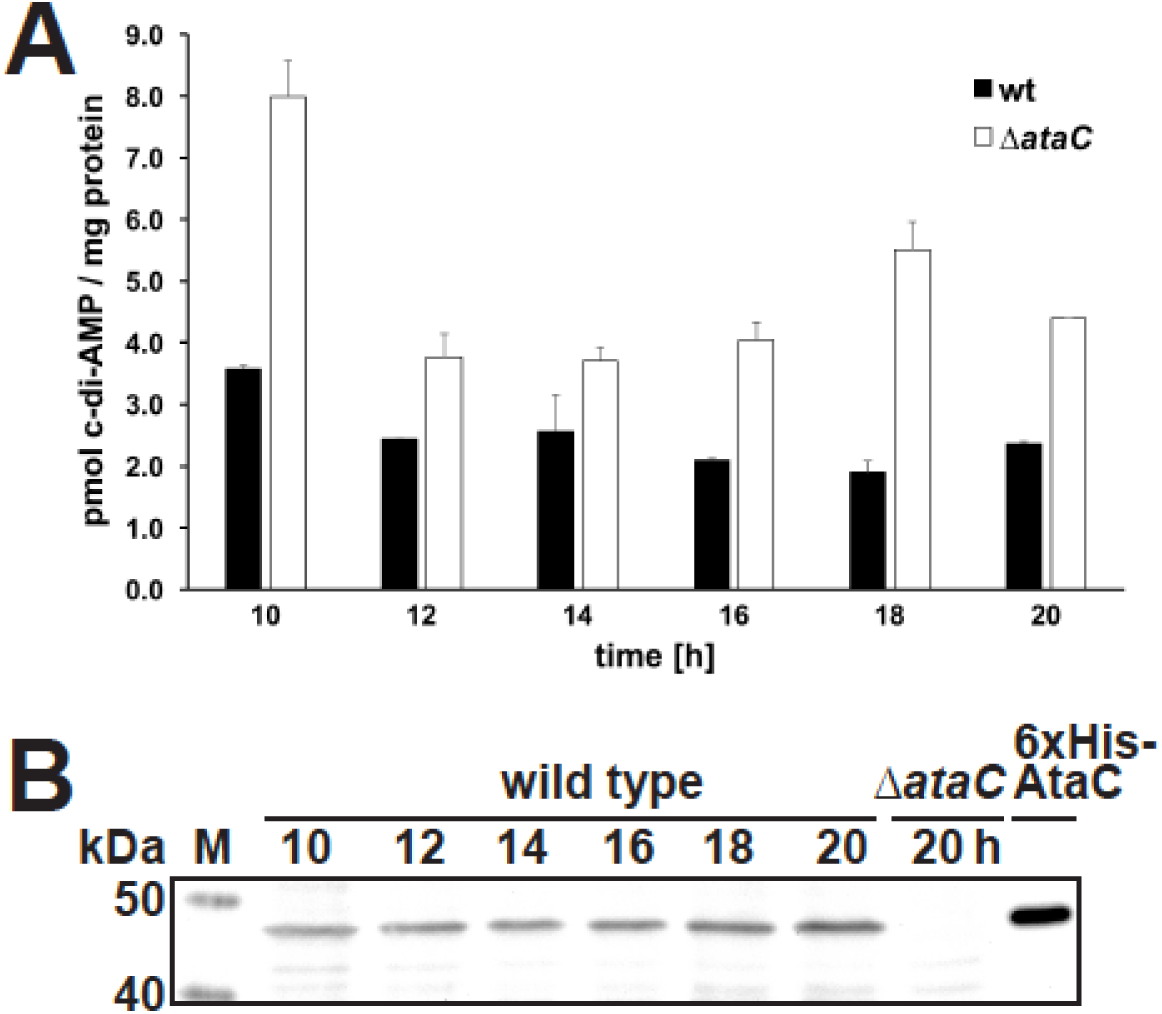
AtaC hydrolyzes c-di-AMP *in vivo* and is constitutively expressed during the life cycle of *S. venezuelae*. (A) Intracellular c-di-AMP levels in *S. venezuelae* wild type and Δ*ataC* during late vegetative growth (10 to 12 h), early sporulation (14 to 16 h) and sporulation (from 18 h). Data are presented as mean of biological replicates ± standard deviation (n=3). (B) Expression profile of AtaC in *S. venezuelae* wild type grown in liquid sporulation medium (MYM). AtaC was detected using a polyclonal anti-AtaC antiserum. Protein samples harvested from Δ*ataC* served as negative control and purified 6xHis-AtaC as positive control, respectively.

### Inactivation of AtaC delays *S. venezuelae* development

To investigate the physiological functions of *disA* and *ataC* and thus of c-di-AMP in *S. venezuelae*, we first analyzed the developmental phenotypes of mutant strains. Colonies of *S. venezuelae* Δ*disA* became green (Figure 5A) and scanning electron microscopy (SEM) confirmed that the Δ*disA* mutant produced spore chains with identical morphology to those of the wild type (Figure 5B). Thus, neither the DisA protein nor the c-di-AMP produced by DisA is required for differentiation.

**Figure 5.**
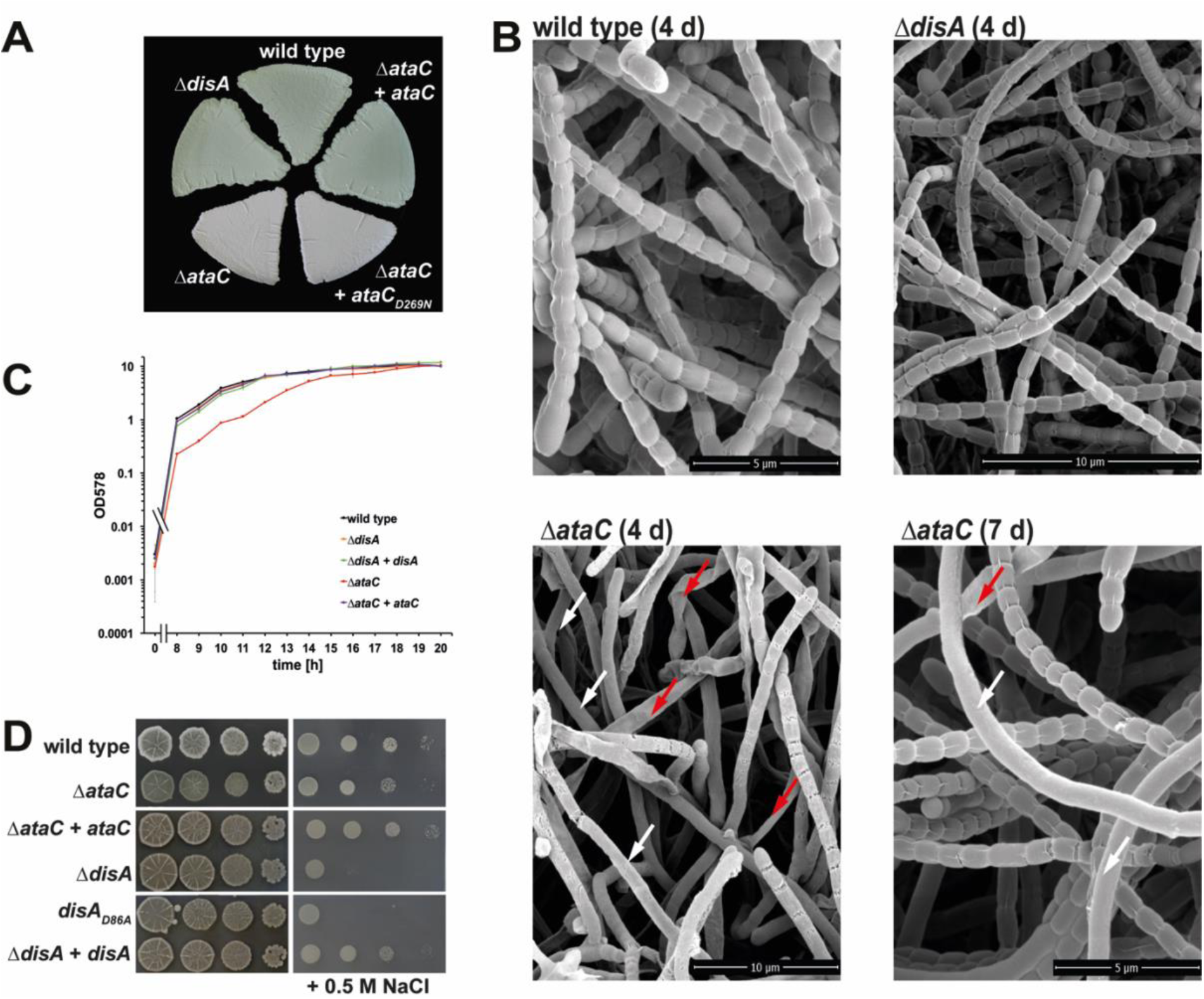
Mutagenesis of c-di-AMP metabolizing enzymes impacts development and ionic stress resistance in *S. venezuelae*. (A) Green morphologies of *S. venezuelae* wild type and Δ*disA* indicate formation of mature spores after 4 days of growth at 30°C on solid sporulation medium (MYM agar). *S. venezuelae* Δ*ataC* fails to accumulate the spore pigment and remains white after the same incubation time. Wild type *ataC* allele complements the Δ*ataC* phenotype but not the enzymatically inactive variant *ataC*_*D269N*_, when expressed *in trans* under control of the native promoter. (B) Scanning electron micrographs showing that after 4 days of incubation on MYM, *S. venezuelae* wild type and Δ*disA* form spores but Δ*ataC* consists predominantly of non-sporulating aerial hyphae (white arrows) and forms flat, likely lysed hyphae (red arrows). After 7 days of growth, Δ*ataC* produced wild type-like spore chains but occasional non-differentiated and lysed hyphae were still detectable. (C) Deletion of *ataC* leads to a growth defect in *S. venezuelae*. c-di-AMP mutants were grown in liquid sporulation medium (MYM) at 30°C and optical density was measured at 578 nm. Δ*ataC* growth is delayed by 3 h and can be restored by expression of the wild type allele under control of its native promoter from the *attB*_*ΦBT1*_ site. (D) Osmotic stress resistance of c-di-AMP mutants. Serial dilutions of spores were spotted on nutrient agar [NA] without additional salt or supplemented with 0.5 M NaCl and grown at 30°C for ∼2 days. Δ*disA* and *disA*_*D86A*_ (expressing inactive DisA) are hypersensitive to salt stress.

In contrast, the *ataC* mutant showed a severe delay in development. After 4 days, the Δ*ataC* strain developed aerial hyphae but did not turn green as the wild type (Figure 5A) and SEM imaging showed mainly undifferentiated aerial hyphae, in contrast to the fully sporulated hyphae seen in the wild type (Figure 5B). Moreover, many of the aerial hyphae of the Δ*ataC* mutant had lysed. After extended incubation (7 days), the aerial hyphae of the Δ*ataC* mutant had largely sporulated, with sporadic non-differentiated and lysed filaments still detected (Figure 5B).

The lysed hyphae seen in the SEMs led us to analyze the growth the Δ*ataC* strain in liquid MYM. As shown in Figure 5C, the *ataC* mutant grew slower than the wild type in exponential phase but reached a similar final OD_578_ after 20 hours. Notably, deletion of *disA* had no effect on growth (Figure 5C).

We could fully complement the defects of Δ*ataC* in development and growth by introduction of the *ataC* wild type allele under the control of its native promoter from the pIJ10170 vector (30) that integrates into the chromosomal *attB*_*ΦBT1*_ site (Figure 5A and S4A). In contrast, expression of the *ataC*_D269N_, which cannot cleave c-di-AMP (Figure S3A) from the same integrative vector did not restore the developmental defects caused by *ataC* deletion (Figure 5A), showing that the cleavage of c-di-AMP by AtaC is crucial for normal development of *Streptomyces*.

Altogether, these results demonstrate that elevated levels of c-di-AMP impair growth and development, whereas reduced levels of c-di-AMP do not affect differentiation under standard growth conditions.

### *disA* mutant is more susceptible to ionic osmostress

Since regulation of osmotic balance is a major function of c-di-AMP in many bacteria (3), we next investigated the osmotic stress resistance of strains with altered c-di-AMP levels due to mutations in either *ataC* or *disA*. We spotted serially diluted spores on nutrient agar (NA) medium plates supplemented with 0.5 M NaCl and a control plate without extra added NaCl. On both plates, the growth of the Δ*ataC* strain was slightly impaired resulting in smaller colony size compared to the wild type (Figure 5D), which likely reflects the growth defect of this strain (Figure 5C). We complemented the growth phenotype of Δ*ataC* with the *ataC* wild type allele expressed *in trans* from the integrative vector pIJ10170 from the *attB*_*ΦBT1*_ site under the control of the native promoter (Figure 5D).

In contrast, when grown on NA plates containing 0.5 M NaCl, Δ*disA* and *disA*_*D86A*_ showed pronounced reduction in growth. Expression of wild type *disA* from pIJ10170 fully complemented the growth defect of Δ*disA* (Figure 5D). The identical Δ*disA* and *disA*_D86A_ phenotypes demonstrate that c-di-AMP produced by DisA is crucial for osmotic stress resistance in *S. venezuelae* (Figure 5D).

In summary, our data revealed that accumulation of c-di-AMP due to *ataC* inactivation, delays development and slows down *Streptomyces* growth in the exponential phase. On the other hand, depletion of c-di-AMP due to *disA* inactivation renders *S. venezuelae* highly susceptible to ionic osmostress.

### The RCK_C domain protein CpeA (Vnz_28055) binds c-di-AMP

RCK_C domains are established direct targets of c-di-AMP that have the I(L)I(L)X_2_DX_1_RX_5_NI(L)I(L) signature for ligand binding (Figure 6A) (31). We found the RCK_C-domain protein Vnz_28055 with a putative c-di-AMP binding motif (Figure 6A-B) in 93 *Streptomyces* species for which complete genome sequences are available (32). We purified N-terminally His-tagged Vnz_28055 and applied differential radial capillary action of ligand assay (DRaCALA) to probe interaction between Vnz_28055 and c-di-AMP. DRaCALA allows visualization of protein-bound radiolabeled ligand as a concentrated ring after the application of the protein-ligand mixture onto nitrocellulose (33). With this assay, we confirmed that Vnz_28055 binds [_32_P]-labeled c-di-AMP (Figure 6C). Excess unlabeled c-di-AMP but not c-di-GMP competed with [_32_P]-c-di-AMP for binding to Vnz_28055. Thus, we identified Vnz_28055 as the first c-di-AMP binding protein in the genus *Streptomyces*.

**Figure 6.**
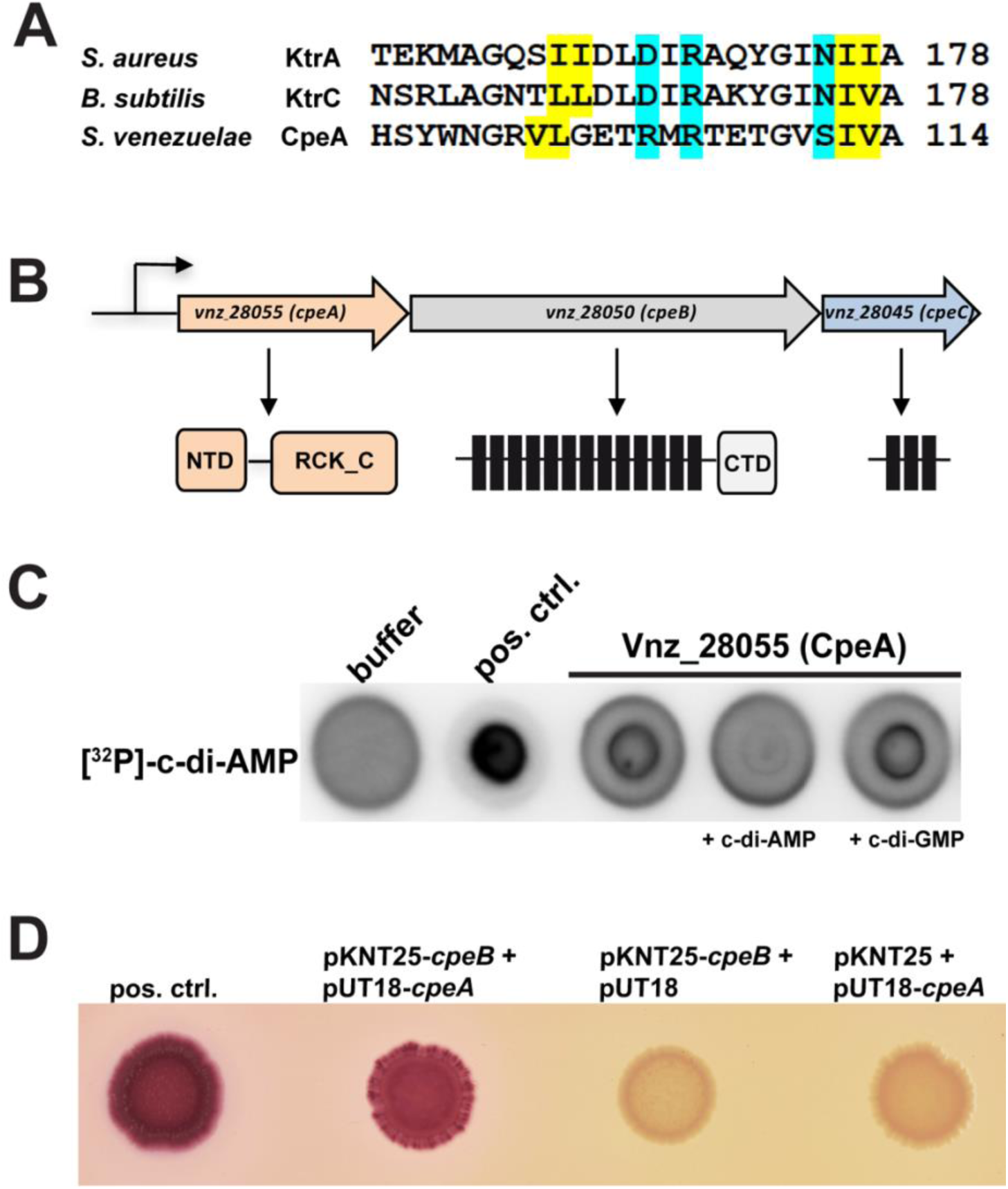
Vnz_28055 (CpeA) binds c-di-AMP. (A) Alignment of the c-di-AMP binding regions in RCK_C (regulator of <u>c</u>onductance of <u>K</u>_+_, <u>C</u>arboxy-terminal) domains was generated using Clustal Omega. C-di-AMP binding residues in KtrA (*S. aureus*;(47)), KtrC (*B. subtilis*;(48)) and conserved amino acids in CpeA are highlighted. Amino acids that form the hydrophobic patch are shown in yellow, residues involved in hydrophilic coordination are highlighted in cyan. (B) *CpeA (vnz_28055), cpeB (vnz_28050)* and *cpeC (vnz_28045)* form an operon in *S. venezuelae*. CpeA has an N-terminal domain (NTD) of unknown function and a C-terminal RCK_C domain. NheB is a predicted structural homolog to the Na_+_/H_+_ antiporter NapA (34). It consists of 13 transmembrane (TM) domains and a cytosolic fraction at the C-terminus (CTD). NheC is a predicted membrane protein with 3 TM domains. (C) CpeA binds [_32_P]-c-di-AMP in DRaCALAs. Binding of the radiolabeled ligand is indicated by dark spots centered on the nitrocellulose. In competition assays, excess (100 µM) cold c-di-AMP or c-di-GMP, respectively, was added to the binding reaction containing [_32_P]-c-di-AMP and 6xHis-CpeA. (D) Adenylate cyclase-based two hybrid assays revealing that CpeA and CpeB interact *in vivo*. Using pKNT25 and pUT18, the T25 and T18 fragments of adenylate cyclase are attached at the C-termini of CpeB and CpeA, respectively. As a positive control, the leucin zipper part of the yeast GCN4 protein was used. Spotted co-transformants were grown for 20 h at 30°C with further incubation at room temperature for ca. 3 days.

*Vnz_28055* forms a conserved operon with *vnz_28050*. Some *Streptomyces* species, such as *S. venezuelae*, contain the small open reading frame *vnz_28045* in the same operon (Figure 6B). Vnz_28050 is a structural homolog of the sodium/proton antiporter NapA (PDB code 5BZ3_A) from *Thermus thermophilus* (34), as predicted with 100 % probability using HHpred (27). To test whether Vnz_28055 and Vnz_28050 form a functional interacting unit, we used a bacterial two-hybrid system in which an interaction between bait and target protein reconstitutes a functional adenylate cyclase (Cya), that allows a *E. coli* Δ*cya* mutant to utilize maltose as a carbon source (35). The two proteins were found to form a complex (Figure 6D), supporting our model that c-di-AMP controls the transport activity of Vnz_28050 by binding to its interaction partner Vnz_28055. Thus, we connect the c-di-AMP function to ionic balance in *Streptomyces* and renamed Vnz_28055-28045 to CpeABC for <u>c</u>ation <u>p</u>roton <u>e</u>xchange component A, B and C.

## DISCUSSION

In this work, using the chloramphenicol-producer *S. venezuelae* as a model and a combination of bioinformatic, biochemical, structural and genetic analyses, we identified AtaC as a novel class of c-di-AMP specific PDEs. AtaC is widely distributed in bacteria and represent the only c-di-AMP PDE in the majority of actinobacteria and an up to now unrecognized c-di-AMP signaling component in pathogens, such as *S. pneumoniae* (Figure 2 and Table S2).

AtaC is a soluble, single-domain phosphodiesterase superfamily protein that is monomeric in solution (Figure S2). In solution, AtaC is structurally similar to the alkaline phosphatase superfamily domain of the C-P bond-cleaving enzyme PhnA from *S. meliloti* 1021 (Figure 3A) (28). As described for DHH-DHHA1 domain-containing proteins GdpP and DhhP, and the HD-domain PDE PgpH, AtaC binds Mn_2+_ to hydrolyze c-di-AMP and we show that residue D269 participates in metal-ion coordination contributing to the active site formation (Figure 3C-E) (17, 22, 36). AtaC has a *k*_cat_ of 0.2 s_-1_ which is comparable to the reported *k*_cat_ of GdpP (0.55 s_-1_). Hydrolytically inactive AtaC_D269N_ has a dissociation constant of 0.7 µM, which is highly similar to the *K*_d_ of wild type PgpH (0.3 - 0.4 µM) (Figures 3G-H) (17, 22). Since we determined the AtaC dissociation constant using a protein carrying the D269N mutation lacking Mn_2+_-coordination, the K_d_ value represents a lower limit as the metal ions bound by the wild type protein likely contribute to c-di-AMP binding. However, while PgpH- and GgdP-type PDEs hydrolyze c-di-AMP exclusively to the linear 5′-pApA, AtaC cleaves c-di-AMP and the intermediate product 5′-pApA to AMP, which has also been shown for DhhP-type PDEs (Figures 2A-B and S1A-B, D) (17, 22, 36). The substrate specificity of AtaC is strictly dependent on two adenosine bases as it shows only weak hydrolysis activity for 5′-pApG and 5′-pGpG in contrast to the DhhP-type PDE TmPDE, which does not distinguish between different nucleobases (Figures 2B and S1E-F) (29).

In *Streptomyces*, AtaC and the DAC DisA are the major regulators of c-di-AMP (Figures 1B and 4A). However, strikingly, the phenotypes of the Δ*ataC* and Δ*disA* mutants with high and low c-di-AMP, respectively, are not invers. On standard growth medium, elevation of intracellular c-di-AMP in Δ*ataC* interferes with growth and ordered hyphae-to-spores transition, while reduction of the second messenger in Δ*disA* does not have any noticeable consequences on these cell functions. On the other hand, when incubated at high external NaCl concentrations, Δ*disA* is severely inhibited in growth, whereas Δ*ataC* grows similarly to the wild type (Figure 5). We found that the RCK_C-domain protein CpeA senses c-di-AMP signals by direct binding of the ligand (Figure 6C). CpeA interacts with CpeB (Figure 6), a structural homolog of the Na_+_/H_+_ antiporter NapA from *T. thermophilus* and a member of the large monovalent cation / proton antiporter (CPA) superfamily (34). Sodium / proton antiporters exist in all living cells, where they regulate intracellular pH, sodium levels, and cell volume (37). In some bacteria, Na_+_/H_+_ antiporters use the proton-motive force to extrude sodium out of the cell and are activated at alkaline pH (38). However, in *Staphylococcus aureus*, the CPA-family transporter CpaA has a cytosolic RCK_C domain that binds c-di-AMP to regulate transport activity (6, 39). Similarly, the regulatory RCK_C-domain proteins KtrA and KtrC bind c-di-AMP to control the activity of the corresponding transport units KtrB and KtrD, respectively (31). Thus, in agreement with this general concept and our data, we propose that c-di-AMP binds to the regulatory RCK_C-domain protein CpeA to activate sodium export via CpeB in *Streptomyces*. At low c-di-AMP, CpeB is presumably inactive allowing accumulation of toxic Na_+_-ions in the cell and leading to growth defects of Δ*disA* on NaCl containing medium. However, on the other hand, likely constant activity of CpeB at high c-di-AMP in Δ*ataC* may result in continuous proton influx affecting intracellular pH and thus important cellular functions causing growth and developmental defects.

In summary, in this study we identified AtaC as a new component of c-di-AMP metabolism in bacteria and uncovered CpeA as the link between c-di-AMP and ion balance in multicellular actinomycetes.

## MATERIAL AND METHODS

For a full explanation of the experimental protocols, see Extended Experimental Procedures in Supplemental Information.

### Bacterial strains and plasmids

All strains, plasmids and oligonucleotides used in this study are listed in Table S1. Plasmids and strains were constructed as described in Extended Experimental Procedures.

### Protein overexpression and purification

*E. coli* BL21 (DE3) pLysS and Rosetta (DE3), respectively, were used for protein overexpression. Cultures were grown in presence of required antibiotics at 37°C and induced with IPTG in the logarithmic phase and transferred for growth at 16°C overnight. Strains overexpressing 6xHis-AtaC, 6xHis-AtaC_D269N_, 6xHis-Vnz_31010, and 6xHis-AtaC_Spn_ were supplemented with MnCl_2_ (17). Cultures were harvested and lysed using a FrenchPress and the proteins were purified via Ni-NTA chromatography. 6xHis-DisA variants and 6xHis-Vnz_28055 were dialyzed twice against 2 L of DisA cyclase buffer (40), and tested PDEs were dialyzed twice against 2 L PDE buffer with 5-10% glycerol (17) at 4°C. Dialyzed proteins were stored at −20 °C. For characterization of biophysical properties of 6xHis-AtaC and 6xHis-AtaC_D269N_, the protein elution was concentrated prior to size exclusion chromatography, flash frozen in liquid nitrogen and stored at −80°C.

### Biochemical characterization of DisA and AtaC variants

Biochemical assays using radioactive-labeled substrates were conducted as described in (32). For diadenylate cyclase (DAC) assays, 5 µM 6xHis-tagged DisA_Sven_, DisA_D86A_ or DisA_Bsu_ were incubated with 83 nM [_32_P]-ATP (Hartmann Analytic) in DisA cyclase buffer. For phosphodiesterase (PDE) assays, 100 nM 6xHis-AtaC or 8 µM 6xHis-Vnz_31010 were mixed with 2 nM [_32_P]-c-di-AMP (Hartmann Analytic, synthesized using purified 6xHis-DisA_Bsu_) in PDE buffer. For competition, 100 µM unlabeled c-di-AMP, c-di-GMP or cAMP were added on ice prior to starting the PDE reactions with [_32_P]-c-di-AMP.

Alternatively, enzymatic activity of 6xHis-AtaC and 6xHis-AtaC_D269N_ was detected by separation of non-labeled reaction products by anion exchange chromatography as described in (29). Reaction solutions contained 50 mM Tris (pH = 7.5), 20 mM NaCl, 100 µM MnCl_2_, 62.5 – 2000 µM ligand (c-di-NMP, 5’-pNpN; N = A or G), 100 nM - 10 µM of 6xHis-AtaC and were incubated at 37°C for 1 h. The reaction was stopped by separating the reaction products from the protein by ultrafiltration (Centricon, 30 kDA cutoff). The filtrate was diluted to 500 µl with running buffer A (50 mM Tris, pH 9) and loaded on a 1 ml Resource™ Q anion exchange column (GE Healthcare Life Sciences). A linear gradient to 40% running buffer B (50 mM Tris, 1 M NaCl, pH 9) over 20 column volumes (CV) was used to separate the nucleotides. The product peaks were identified by comparison to nucleotide standards, c-di-NMP, pNpN, N = A or G obtained from BioLog.

### Differential radial capillary action of ligand assay

DRaCALAs were performed using 2 µg of purified 6xHis-CpeA (Vnz_28055) as described in Roelofs et al 2011 (33) with minor modifications. Purified HD domain of PgpH from *L. monocytogenes* (17) fused to an N-terminal GST tag was used as a positive control. For competition, reactions were supplemented 100 µM of non-labeled c-di-AMP or c-di-GMP prior to addition of [_32_P]-c-di-AMP.

### Western blotting

For detection of 3xFLAG-tagged DisA, Western blot analysis was performed as described in (32) using 5 µg total protein of *S. venezuelae* Δ*disA* expressing the FLAG-tagged *disA* allele from the *Φ*_*BT1*_ integration site under the control of the native promoter. Anti-FLAG primary antibody (Sigma) and the anti-mouse IgG-HRP (Thermo Fisher Scientific) were used for detection. AtaC was detected in the wild type strain (10 µg total protein) using polyclonal rabbit anti-AtaC antiserum as primary antibody (generated by Pineda GmbH using purified 6xHis-AtaC) and donkey anti-rabbit-HRP secondary antibody (GE Healthcare). ECL chemiluminescent detection reagent (Perkin Elmer) was used for visualization.

### c-di-AMP extraction and quantification

The nucleotide extraction protocol from (2) was adapted to *Streptomyces*. Wild type, Δ*disA* and Δ*ataC* strains were grown in MYM. Samples for c-di-AMP extraction and for determination of the protein concentration were taken every 2 h after initial growth for 10 h. c-di-AMP was extracted using acetonitrile/methanol from cells disrupted using the BeadBlaster (Biozym). Samples were analyzed using LC-MS/MS as described in (2).

### Bacterial Adenylate Cyclase Two-Hybrid (BACTH) assays

BACTH system was used to assay protein-protein interaction of CpeA and CpeB *in vivo* (35). Plasmids expressing C-terminal fusions of CpeA and CpeB to T18 and T25 fragments of *cyaA* from *Bordetella pertussis*, respectively, were transformed into *E. coli* W3110 lacking *cya* (41). Co-transformants were spotted on MacConkey agar supplemented with maltose (1%), ampicillin (100 µg/ml), and kanamycin (50 µg/ml). Red colonies indicate cAMP-dependent fermentation of maltose which occurs upon direct interactions of the proteins fused to the otherwise separate adenylate cyclase domains.

### Small-angle X-ray scattering

Size-exclusion chromatography coupled small-angle X-ray scattering data (42, 43) for AtaC were collected at the EMBL Hamburg P12 beamline at PETRA3 (DESY, Hamburg). CHROMIXS of the ATSAS Suite (44) was used for analysis and processing of the chromatogram results. In brief, after choosing an appropriate buffer region and averaging of the respective frames, the protein scattering frames from the elution peak were buffer subtracted and averaged. The final protein scattering data were then analyzed using the ATSAS suite. The theoretical scattering curve of the AtaC model derived from HHpred/MODELLER was obtained using CRYSOL (45). Ab initio models were calculated using DAMMIF and averaged using DAMAVER as described earlier (29).

### Nano differential scanning fluorimetry

Thermal unfolding experiments of AtaC were performed with a Tycho NT.6 instrument (NanoTemper Technologies). The samples were heated in a glass capillary at a rate of 30 K/min and the internal fluorescence at 330 nm and 350 nm was recorded. Data analysis, data smoothing and calculation of derivatives was done using the internal evaluation features of the Tycho instrument.

### Bioinformatic characterization of AtaC and its abundance in prokaryotes

AtaC was identified as a member of the phosphodiesterase family of proteins by annotation of the *S. venezuelae* genome with interproscan (version 5.27-66.0; http://dx.doi.org/10.7717/peerj.167) and searching for proteins harboring type I phosphodiesterase / nucleotide pyrophosphatase domain (Pfam: PF01663).

### Scanning electron microscopy (SEM)

SEM was performed as previously described (46).

## ACKNOWLEDGEMENTS

We are grateful to Mark J. Buttner and Fabian M. Commichau for helpful discussion and critical reading of the manuscript and thank Matt Bush for technical support with scanning electron micrographs. We thank the staff of the EMBL-Hamburg beamline P12 at PETRA3 (EMBL/DESY, Hamburg, Germany) for outstanding scientific support. We also acknowledge Anna-Lena Hagemann and Annette Garbe for technical support with LC-MS/MS funded by the DFG Priority Program SPP 1879 (KA 730/9-1). Research in Gregor Witte’s lab is funded by DFG GRK1721 and the DFG Priority Program SPP 1879 (WI 3717/3-1). Research in Natalia Tschowri’s lab is funded by the DFG Emmy Noether-Program (TS 325/1-1) and the DFG Priority Program SPP 1879 (TS 325/2-1).

## AUTHOR CONTRIBUTIONS

N.T. designed the study. All authors designed and interpreted experiments, which were performed by A.L., D.J.D., M.M.A-B, G.W., V.K. and K.C.F. The figures were made by A.L., D.J.D., M.M.A-B, G.W. and N.T. The paper was written by A.L., D.J.D., G.W. and N.T. with input from the other authors.

