## Extended Material and Methods and Supplemental Data for "c-di-AMP hydrolysis by a novel type of phosphodiesterase promotes differentiation of multicellular bacteria"

### SUPPLEMENTARY EXPERIMENTAL PROCEDURES

#### Bacterial strains and growth conditions

All strains used in this study are listed in Table S1. *Escherichia coli* strains were grown in LB medium under aerobic conditions at 37°C. If required, LB was supplemented with 100 µg/ml ampicillin (Amp), 50 µg/ml kanamycin (Kan), 50 µg/ml apramycin (Apr) and/or 15 µg/ml chloramphenicol (Cam). If hygromycin B (Hyg) was used, LB agar was replaced by Nutrient Agar (NA; Roth) and LB was substituted by LBon (LB without salt) with addition of 16 µg/ml and 22 µg/ml Hyg, respectively. *S. venezuelae* strains (Table S1) were grown aerobically at 30 °C in liquid Maltose-Yeast Extract-Malt Extract (MYM) medium (1) supplemented with trace element solution (2) or on MYM agar. For growth analysis, 50 ml MYM were inoculated with spores at a final concentration of $10^6$ CFU/µl and OD was measured at 578 nm. To study development, 12 µl of $10^5$ CFU/ml *S. venezuelae* spores were spread as patches on MYM agar and bacteria were photographed using a Canon EOS 1300D (W) camera after 4 days of growth at 30 °C. For osmostress experiments, 10 µl of serially diluted of *S. venezuelae* spores ($10^1$ to $10^4$ CFU/µl) were dropped on NA medium with or without 0.5 M NaCl, respectively. Plates were incubated at 30°C and pictures were taken using a Canon EOS 1300D (W) camera. For *in vivo* interaction studies using Bacterial Adenylate Cyclase Two-Hybrid (BACTH) assays three corresponding single transformants of *E. coli* W3110 Δ*cya* were suspended in 1 ml of sterile phosphate buffered saline (PBS) and 3 µl of the resulting suspension was spotted on a plate containing MacConkey Base Agar (Difco) supplemented with Amp (100 µg/ml), Kan (50 µg/ml) and maltose (1%). The plate was photographed using a Canon EOS 1300D (W) camera after growth at 30 °C for 20 h with further incubation at room temperature for three days.

**Generation and complementation of *S. venezuelae* c-di-AMP mutants**

Oligonucleotides and bacterial strains used for mutagenesis are listed Table S1.

*Generation of vnz_27310 (ataC) deletion mutant and phage transduction.* The *ataC* deletion was conducted using a modified Redirect PCR targeting protocol (3, 4). *E. coli* BW25113/pIJ790 cells induced for λ Red mediated recombination and containing cosmid PI1_F15 (with *ataC*) were transformed with PCR amplified *aac(3)IV-oriT* (*apr-oriT*) cassette extended with regions homologous to the *ataC* locus. After growth of transformants in presence of apramycin only, cosmids were isolated and re-transformed into *E. coli* W3110 with selection for Apr resistance. Correct integration of the Apr resistance cassette was verified by PCR using test primers annealing to the *ataC* flanking region (Table S1). PI1_F15 *ataC::apr* was transformed into strain ET12567/pUZ8002 and conjugated into wild type *S. venezuelae.* Bacteria were plated on SFM agar, incubated overnight at room temperature (RT), overlaid with 20 µl of 25 mg/ml nalidixic acid (Nal) and 50 mg/ml Apr in 2 ml ddH$_2$O and incubated at 30°C. *S. venezuelae* colonies were selected on NA containing Apr and Nal to remove *E. coli*. Kan$^S$ and Apr$^R$ mutants were confirmed by PCR.

The *ataC::apr* allele was transduced into a new wild type background via SV1 phage using a modified protocol from (5). Briefly, SV1 wild type phages were diluted and mixed with *S. venezuelae* Δ*ataC* spores. After overnight incubation at 30°C, plates were soaked with LB at RT and phages were gathered and filtered using a 0.45 µm filter. 100 µl phages containing the *ataC::apr* allele were plated with wild type spores on MYM agar and incubated overnight at RT. For selection of desired transductants, plates were overlaid with 20 µl 50 mg/ml Apr in 2 ml ddH$_2$O. The *ataC* deletion was confirmed by PCR.

*Generation of disA deletion mutant*. The *disA* mutant was generated by transduction of the *disA::apr* allele (6) into the *S. venezuelae* wild type using SV1 phage.

*Generation of disA*$_{D86A}$ *point mutation.* The point mutation was introduced using a combination of modified Redirect PCR targeting and single stranded DNA recombineering protocols as described in (7).The Kan resistance cassette in cosmid SV-4-B12 (containing *disA*) was exchanged with *apr-oriT* extended with homologous regions to *neo*. The resulting cosmid was used to generate the *E. coli* strain HME68/SV-4-B12 *neo::apr-oriT* which was induced for λ Red recombination and electroporated with the mutagenic oligonucleotide disA_D86Achr_rev and oligo100 (8) and plated on MacConkey agar containing 1% galactose and Apr. Red, Apr$^{R}$ clones were analyzed for the *disA::disA*$_{D86A}$ allele by PCR using a primer pair specific for the D86A mutation and sequencing. Purified cosmid was electroporated into ET12567/pUZ8002 and conjugated into wild type *S. venezuelae*. Single colonies were selected on NA medium containing Apr and Nal followed by growth on NA without antibiotics to achieve loss of the cosmid. Colonies sensitive to Apr were verified for the *disA* D86A mutation by sequencing.

*Complementation of the deletion mutants.* To complement the *disA* deletion, a DNA fragment containing the wild type allele with 528 bp upstream of the start codon (including *disA* promoter) was cloned into pIJ10170 or its derivative p3xFLAG which allow integration at the *attB*$_{\Phi BT1}$ site in the *S. venezuelae* chromosome. pIJ10170-*disA* and p3xFLAG-*disA*, respectively, were conjugated into *disA::apr* using *E. coli* ET12567/pUZ8002. After selection on NA medium containing Nal and Hyg, Hyg$^{R}$ colonies were grown on MYM agar for spore stock generation.

The complementation of the *ataC* deletion mutant with the wild type allele was conducted similarly. Here, using an overlap PCR, a DNA fragment corresponding to 200 bp upstream of the start codon of *vnz_27305* was fused to full length *ataC* resulting in the construct *vnz_27305prom-ataC*. The fragment was cloned into pIJ10170 resulting in the plasmid pSVAL11 and introduced into the Δ*ataC* chromosome as described above. For complementation of Δ*ataC* with the D269N allele, pSVAL11 was used as template in a backbone PCR to amplify a circular construct with the primers D269N_backbone_f and D269N_backbone_r. The plasmid pIJ10170-*vnz_27305prom-ataC*$_{D269N}$ was generated, the mutation confirmed by sequencing and conjugated into Δ*ataC* as described above.

**Construction of plasmids**

Oligonucleotides used for cloning are listed in Table S1. *disA*, *vnz_31010*, *ataC* and *vnz_28055* were amplified from *S. venezuelae* genomic DNA (gDNA). D86A point mutation in *disA* was introduced by following the four-primer/two-step PCR protocol (9). PCR products of all constructs were cloned into the pET15b vector. The *ataC*$_{\mathrm{D269N}}$ construct was obtained using quick change site directed mutagenesis using the pET15b-*ataC* plasmid as a template. Codon optimized *ataC*$_{Spn}$ was synthesized *de novo* and cloned into pET15b via the NdeI and BamHI restriction sites by GenScript.

Full-length *cpeA* (*vnz_28055*) and *cpeB* (*vnz_28050*) excluding the respective stop codons were cloned into pUT18 and pKNT25 (Euromedex). The resulting constructs carry in-frame fusions of the sequences encoding the T18 and T25 fragments of *cyaA* from *Bordetella pertussis* to the 3’ end of *cpeA* and *cpeB*, respectively. Expression of the fused genes is under control of the *lac* promoter.

**Protein overexpression and purification**

pET15b constructs were transformed into *E. coli* BL21 (DE3) pLysS. Strain Rosetta (DE3) pET28-$disA_{Bsu}$ was directly used for overexpression. 1 L LB containing Amp and Cam (and 0.2% glucose in case of PDEs) was inoculated 1:100 with overnight cultures and grown with shaking at 37°C. For $DisA_{Bsu}$ overexpression Amp was replaced with 25 µg/ml Kan. Cultures were induced with a final concentration of 0.1-0.2 mM IPTG at $OD_{578}$ between 0.5 and 0.7; cultures for PDE overexpression were supplemented with 0.35 mM $MnCl_2$ (10). Proteins were overexpressed overnight at 16°C and shaking. Subsequently, cultures were pelleted and lysed using a FrenchPress. Strains expressing DisA variants and 6xHis-Vnz_28055 were lysed in DisA lysis buffer (20 mM Tris HCl, pH 8; 300 mM NaCl, 10% glycerol, 20 mM imidazole; 0.05% Triton X-100; 0.5 mM DTT; 5 mM $MgCl_2$) supplemented with cOmplete protease inhibitor cocktail tablets, EDTA-free (Roche). Strains expressing 6xHis-Vnz_31010, 6xHis-AtaC and 6xHis-$AtaC_{Spn}$ were lysed in PDE lysis buffer containing cOmplete protease inhibitor cocktail tablets, EDTA-free (similar to DisA lysis buffer but Tris HCl, pH 8 replaced by 20 mM Tris HCl 7.5; $MgCl_2$ replaced by 10 mM $MnCl_2$). Clarified lysate supernatants of 6xHis-tagged proteins were loaded on 0.5-1 ml 50% Ni-NTA SuperFlow (iba) overnight at 4°C. Then, the matrix was washed with respective lysis buffers. DisA protein variants and 6xHis-Vnz_28055 were eluted with the following buffer: 50 mM Tris HCl, pH 8; 300 mM NaCl; 10% glycerol; 250 mM imidazole; 0.5 mM DTT; 5 mM $MgCl_2$. The PDE elution buffer was similar to DisA elution buffer but containing 50 mM Tris HCl, 7.5 instead of Tris HCl, pH 8 and 10 mM $MnCl_2$ instead of $MgCl_2$. Fractions containing eluted proteins (identified by Coomassie staining of 12% polyacrylamide gels) were pooled. Eluates of DisA variants and 6xHis-Vnz_28055 were dialyzed twice against 2 L of DisA cyclase buffer (25 mM Tris HCl, pH 8; 250 mM NaCl, 10 mM MgCl, 5 mM β-mercaptoethanol, 10% glycerol (modified from (11), and tested PDEs were dialyzed twice against 2 L PDE buffer with 5-10% glycerol (20 mM Tris HCl, pH 7.5; 50 mM NaCl; 10 mM $MnCl_2$ (modified from(10) at 4°C under stirring. Dialyzed proteins were stored at -20°C until further use in diadenylate cyclase (DAC), differential radial capillary action of ligand (DRACaLA) or phosphodiesterase (PDE) assays.

For characterization of biophysical properties of 6xHis-AtaC and 6xHis-AtaC$_{D269N}$, Rosetta (DE3) cell pellets containing pET15b-*ataC* and pET15b-*ataC*$_{D269N}$ constructs, respectively, were resuspended in buffer A (20 mM HEPES, 300 mM NaCl, 20 mM Imidazole, 10% glycerol, 0.5 mM $MnCl_2$, pH 7.5) and lyzed by sonication. After centrifugation, clear supernatant was loaded on Ni-NTA columns. The columns were washed with buffer A and the protein was eluted with buffer B (20 mM HEPES, 100 mM NaCl, 250 mM imidazole, 10% glycerol, 0.5 mM $MnCl_2$, pH 7.5). The protein elution was concentrated prior to size exclusion chromatography on a HiLoad Superdex 200 column (GE Healthcare) equilibrated with buffer C (20 mM HEPES, 100 mM NaCl, 0.5 mM $MnCl_2$, pH 7.5). The pure protein was concentrated, flash frozen in liquid nitrogen and stored at -80°C.

**c-di-AMP extraction and quantification**

The nucleotide extraction protocol from (12) was adapted to *Streptomyces*. Wild type, Δ*disA* and Δ*ataC* strains were grown in 100 ml MYM. Beginning with 10 h, 5 ml samples for c-di-AMP extraction and two 1 ml samples for protein determination were taken every 2 h. c-di-AMP samples were centrifuged at 4000 rpm and 4°C for 15 min using a swing rotor (Heraeus Megafuge 16R, Thermo Scientific), frozen in liquid nitrogen and stored at -80°C. Protein samples were centrifuged at max. speed and stored at -20°C.

For c-di-AMP extraction, samples were suspended in 800 µl Extraction mixture II (acetonitrile/methanol/water [2:2:1]), transferred into 2 ml screw cap tubes prefilled with 0.1 mm silica beads (Biozym), shock frozen for 15 s in liquid nitrogen and heated for 10 min at 95°C. After cooling on ice, samples were disrupted using the BeadBlaster at 4°C with 2 cycles at 6 m/s for 45 s and 2 min interval. Samples were cooled for 15 min on ice and centrifuged at max. speed and 4°C for 15 min. Supernatants were transferred into a 2 ml reaction tubes. Remaining pellets were suspended in 600 µl Extraction mixture I (acetonitrile/methanol [1:1]), pulsed two times for 30 s at 6 m/s with a 60s interval, incubated on ice and centrifuged as above. The extraction with 600 µl Extraction mixture I was repeated once. All supernatants (~2 ml) were combined and stored for protein precipitation for two days at -20°C. Precipitated proteins were removed by centrifugation and the precipitation step was repeated. Finally, samples were air dried in a SpeedVac Plus SC110A connected to Refrigerated Vapor Trap RVT100 (Thermo Scientific) at low temperature settings and analyzed using LC-MS/MS as described in (12).

Samples for protein quantification were suspended in 800 µl 0.1 M NaOH, transferred into 2 ml screw cap tubes prefilled with 0.1 mm silica beads (Biozym) and heated for 10 min at 98°C. Cell lysis was performed in BeatBlaster with 2 pulses for 30 s at 6 m/s and an interval of 2 min. Lysates were centrifuged at max. speed and 4°C for 15 min. Supernatant was saved and the extraction step was repeated. Supernatants were combined and protein concentration was determined via Bradford using Roti-Quant.

For normalization of c-di-AMP concentration to the protein amount, following formula was used:

$$\frac{c{-}di{-}AMP\ [nM] \cdot 200}{cV\ [ml] \cdot c590\ [\frac{\mu g}{ml\ cells}]} = \frac{c{-}di{-}AMP\ [pmol]}{protein\ [mg]}$$

**Isothermal Titration Calorimetry**

ITC experiments were performed using a Malvern PEAQ-ITC system with 21 µM protein in ITC buffer (20 mM HEPES, pH = 7.5; 100 mM NaCl) in the cell. The respective nucleotides (210 µM) were titrated into the cell by 19 injections of 2 µl, spaced 150 s apart, at 25°C. The data was analyzed using the MicroCal PEAQ-ITC analysis software provided with the instrument. All titrations were repeated to confirm robustness of the assay.

**Size-exclusion coupled static light scattering**

Determination of the molecular weight of AtaC was performed using a 24 ml Superdex S200 increase size-exclusion coupled to multi-angle laser light scattering and refractive index monitor (WYATT miniDAWN TREOS, WYATT Optilab T-rEX). Data were analyzed using the ASTRA software package provided with the instrument (Wyatt).

**Bioinformatic characterization of AtaC and its abundance in prokaryotes**

A local PATRIC database was installed and used to determine the conservation of AtaC (PGF_00172869) across prokaryotes (Suppl. Table 2). Duplicated entries with identical genome were removed (177 in total), but keeping the first entry. The same database was also used to determine the conservation of DisA (PGF_00421347), PgpH homologues (PGF_03110657), GdpP-type proteins (PGF_00033444) and DhhP-like proteins (PGF_01833449). An in-house python script was used to extract taxonomic information for each of the AtaC homologues. Specifically, the accession number of each species was used to access NCBI taxonomy and the taxonomic information was integrated with the original PATRIC table. Only phyla with more than 5 genomes were kept, and entries with no taxonomic information were excluded from the analysis. Finally, a total of 5200 entries were used to generate the AtaC abundance pie chart. Multiple sequence alignment of AtaC proteins (see Suppl. Table S2 for respective PATRIC IDs) from different phyla was performed using CLUSTAL OMEGA (1.2.4) on https://www.ebi.ac.uk/Tools/msa/clustalo/.

### SUPPLEMENTARY INFORMATION

**Table S1. Strains, plasmids and oligonucleotides used in this study**

| | **Genotype or comments** | **Source or reference** |
|---|---|---|
| **Strains** | | |
| ***S. venezuelae*** | | |
| NRRL B-65442 | Wild type | (NCBI Reference Sequence: NZ_CP018074.1) |
| *disA::apr* | ATCC 10712 *SVEN_3211::aac(3)IV*; Apr$^{R}$ | (6) |
| SVAL5 | Δ*disA::apr*, *attB*$_{\Phi BT1}$::p3xFLAG-*disA*; Apr$^{R}$, Hyg$^{R}$ | This study |
| SVAL8 | *disA::disA*$_{D86A}$ | This study |
| SVAL19 | Δ*disA* (SV1-transduction); Apr$^{R}$ | This study |
| SVAL20 | Δ*vnz_27310::apr* (Redirect); Apr$^{R}$ | This study |
| SVAL22 | Δ*ataC::apr* (SV1-transduction from SVAL20); Apr$^{R}$ | This study |
| SVAL24 | Δ*disA::apr*, *attB*$_{\Phi BT1}$::pIJ10170-*disA*; Apr$^{R}$, Hyg$^{R}$ | This study |
| SVAL26 | Δ*ataC::apr*, *attB*$_{\Phi BT1}$::pIJ10170-*vnz_27305prom-ataC*; Apr$^{R}$, Hyg$^{R}$ | This study |
| SVAL27 | Δ*ataC::apr*, *attB*$_{\Phi BT1}$::pIJ10170-*vnz_27305prom-ataC*$_{D269N}$; Apr$^{R}$, Hyg$^{R}$ | This study |
| ***E. coli*** | | |

| | | |
|---|---|---|
| W3110 | K-12 derivative; *F-, λ- ,rpoS(Am), rph-1, Inv(rrnD-rrnE)* | (13) |
| W3110 Δ*cya* | W3110 derivative with deleted adenylate cyclase | (14) |
| ET12567/pUZ8002 | *dam, dcm, hsd*; $Kan^R$, $Cm^R$ | (15) |
| BW25113/pIJ790 | (Δ(*araD-araB*)*567*, Δ*lacZ4787*(::*rrnB-4*), *lacIp-4000*($lacI^Q$), λ-, *rpoS369*(Am), *rph-1*, Δ(*rhaD-rhaB*)568, *hsdR514*; $Cm^R$ | (16) |
| BL21 (DE3) pLysS | F– *ompT hsdS*(rB– mB–) *gal dcm* λ(DE3), $Cm^R$ | Promega |
| HME68 | W3110 Δ(*argF-lac*)*U169 galKtyr145UAG mutS<>cat* | (17) |
| Rosetta (DE3) pET28-*disA*$_{Bsu}$ | Overexpression of *Bacillus subtilis* DisA; $Kan^R$, $Cm^R$ | (18) |
| Rosetta 2 (DE3) | $F^-$ *ompT hsdS*$_B$($r_B^-$ $m_B^-$) *gal dcm* (DE3) pRARE2 ($Cam^R$) | Novagen |
| **Plasmids** | | |
| pIJ773 | Plasmid template for amplification of the *apr-oriT* cassette for 'Redirect' PCR-targeting; $Apr^R$ | (3) |
| pIJ790 | Modified λRED recombination plasmid [*oriR101*] [*repA101*(ts)] *araBp-gam-be-exo*; $Cm^R$ | (3) |
| pIJ10170 | pMS82 derivative; $Hyg^R$ | (19) |
| pUZ8002 | RP4 derivative with defective oriT; $Kan^R$ | (15) |
| p3xFLAG | pIJ10170 derivative containing *3xFLAG* sequence downstream of MCS; $Hyg^R$ | (20) |
| pGEX_6P_1 | T7 expression vector (modified MCS); $Amp^R$ | GE Healthcare |
| pET15b | T7 expression vector; $Amp^R$ | Novagen |
| pKNT25 | Low copy vector encoding the T25 fragment of *Bordetella pertussis cyaA* downstream of the MCS; $Kan^R$ | Euromedex |

| | | |
|---|---|---|
| pUT18 | High copy vector encoding the T18 fragment of *B. pertussis cyaA* downstream of the MCS; Amp$^R$ | Euromedex |
| pECAL1 | pET15b-*disA*; Amp$^R$ | This study |
| pECAL4 | pET15b-*disA*$_{D86A}$; Amp$^R$ | This study |
| pECAL12 | pET15b-*ataC*; Amp$^R$ | This study |
| pECAL13 | pET15b-*vnz_31010*; Amp$^R$ | This study |
| pECAL16 | pET15b-*ataC*$_{Spn}$; Amp$^R$ | This study |
| pECAL17 | pET15b-*vnz_28055 (cpeA)*; Amp$^R$ | This study |
| pECAL18 | pKNT25-*vnz_28050 (cpeB)*; Kan$^R$ | This study |
| pECAL19 | pUT18-*vnz_28055 (cpeA)*; Amp$^R$ | This study |
| pSVAL6 | pIJ10170-*disA*; Hyg$^R$ | This study |
| pSVAL11 | pIJ10170-*vnz_27305prom-ataC* (200 bp upstream of *vnz_27305* start codon fused to *ataC*); Hyg$^R$ | This study |
| pSVAL12 | pIJ10170-*vnz_27305prom-ataC*$_{D269N}$ (pSVAL11 derivative); Hyg$^R$ | This study |
| pSVNT-10 | p3xFLAG-*disA*; Hyg$^R$ | This study |
| pDD29 | pET15b-*ataC*$_{D269N}$ (pECAL12 derivative); Amp$^R$ | This study |

Underlined nucleotides indicate restriction sites, nucleotides in bold represent introduced mutations and nucleotides in italics indicate sequences overlapping to other genes

| Oligonucleotide | Sequence |
|---|---|
| **Oligonucleotides used for chromosomal *disA* D86A point mutation, PCR verification and sequencing** | |
| disA_D86Achr_rev | TCTTGGTGATGTCCTTGTCGAGGACGAGCGCGCC**CG**CGAGCTTGCACAGCTCCCGCAGCCGCGTGGCGGC |
| disA_D86A_check_fwd | GGAGCTGTGCAAGCTCGCG |
| disA_fwd_NdeI | TATCATATGGTGGCAGCCAAGGAC |

| | |
|---|---|
| disA_rev_XhoI | TATCTCGAGCTAGACGTACCGCTCAAG |
| **Oligonucleotides used for verification of *disA* deletion** | |
| disA_test_f | GTGGTTCACTCACGCCGCATGAACGGTTC |
| disA_test_r | GGCACGTACCTGGTGGAGGCGAAGGTG |
| **Oligonucleotides used for complementation of *disA* deletion with *disA-3xFLAG*** | |
| 3142_NdeI-for | GCTGCATATGGGCCGGCGGGTCG |
| 3142_XhoI-rev | GCAGCCTCGAGGACGTACCGCTCAAGGATC |
| **Oligonucleotides used for complementation of *disA* deletion with wild type *disA*** | |
| 3142_NdeI-for | GCTGCATATGGGCCGGCGGGTCG |
| disA_rev_XhoI | TATCTCGAGCTAGACGTACCGCTCAAG |
| **Oligonucleotides used for *vnz_27310 (ataC)* deletion and PCR verification** | |
| 27310_fwd_Apra | CGAAGCGATCGCGGCCACCGCCGCGCCCACCCGCTGATG*ATTCCGGG GATCCGTCGACC* |
| 27310_rev_Apra | GGTCGTGGGGGGGGAGGAGACGGGTGAGGAGTGGGCTCA*TGTAGGC TGGAGCTGCTTC* |
| 27310_test_f | ACACCGTGCGGCAGACCC |
| 27310_test_r | TTCCCGCAGTCCATGGTTCC |
| **Oligonucleotides used for complementation of *ataC* deletion (sequences overlapping to putative promoter in italics)** | |
| 27305prom_f_NdeI | TATCATATGGGTGTCCCGGCTCGTCGAC |
| 27305prom_r_OL_ataC | GCTGCACCAT*GGACTCCATCCTACGGGGCT* |
| ataC_f_OL_27305prom | *GATGGAGTCC*ATGGTGCAGCCGACCGCCGT |
| 5409_rev_XhoI | TATACTCGAGTCAGGTGCGGACTTCGAG |
| **Oligonucleotides used for generation of *pIJ10770-27305prom-ataC*$_{D269N}$** | |

| | |
|---|---|
| D269N_backbone_f | CGGCGCTGTACGTCACGGCC**A**ACCACGGCATGGTCGACAT |
| D269N_backbone_r | ATGTCGACCATGCCGTGGT**T**GGCCGTGACGTACAGCGCCG |
| **Oligonucleotides used for generation of pET15b overexpression constructs** | |
| disA_fwd_NdeI | TATCATATGGTGGCAGCCAAGGAC |
| disA_rev_XhoI | TATCTCGAGCTAGACGTACCGCTCAAG |
| disA_D86A_fwd | GCAAGCTCG**CG**GGCGCGCTC |
| disA_D86A_rev | GAGCGCGCC**CG**CGAGCTTGC |
| 5409_fwd_NdeI | TATCATATGATGGTGCAGCCGACCG |
| 5409_rev_XhoI | TATACTCGAGTCAGGTGCGGACTTCGAG |
| 6143_fwd_NdeI (for *vnz_31010*) | TATCATATGGTGATCGTCATCGCCCATGT |
| 6143_rev_BamHI (for *vnz_31010*) | TATGGATCCTCAGACCGGCACGGTC |
| 28055_fwd_NdeI | TAT CATATG GTGCCTGCTCCACGGATG |
| 28055_rev_XhoI | TATA CTCGAG TCACTCCCGTCCGAGTATGG |
| **Oligonucleotides used for generation of the pET15b-*ataC*$_{D269N}$ construct** | |
| DD60AtaC(D269N)_fwd | GTACGTCACGGCC**A**ACCACGGCATGGTCGA |
| DD61AtaC(D269N)_rev | GGCCGTGACGTACAGCGCCGAGC |
| **Oligonucleotides used for generation of the constructs for interaction studies** | |
| 28050_NT25f_XbaI | TATTCTAGACGTGCATTCCGCTCTGTTCCT |
| 28050_NT25r_KpnI | TATGGTACCCGCGCGCGGCCGGCCGGATCCT |
| 28055_T18f_KpnI | TATGGTACCGCCTGCTCCACGGATGAGC |
| 28055_T18r_EcoRI | TAGCAGAATTCGACTCCCGTCCGAGTATGGAGG |

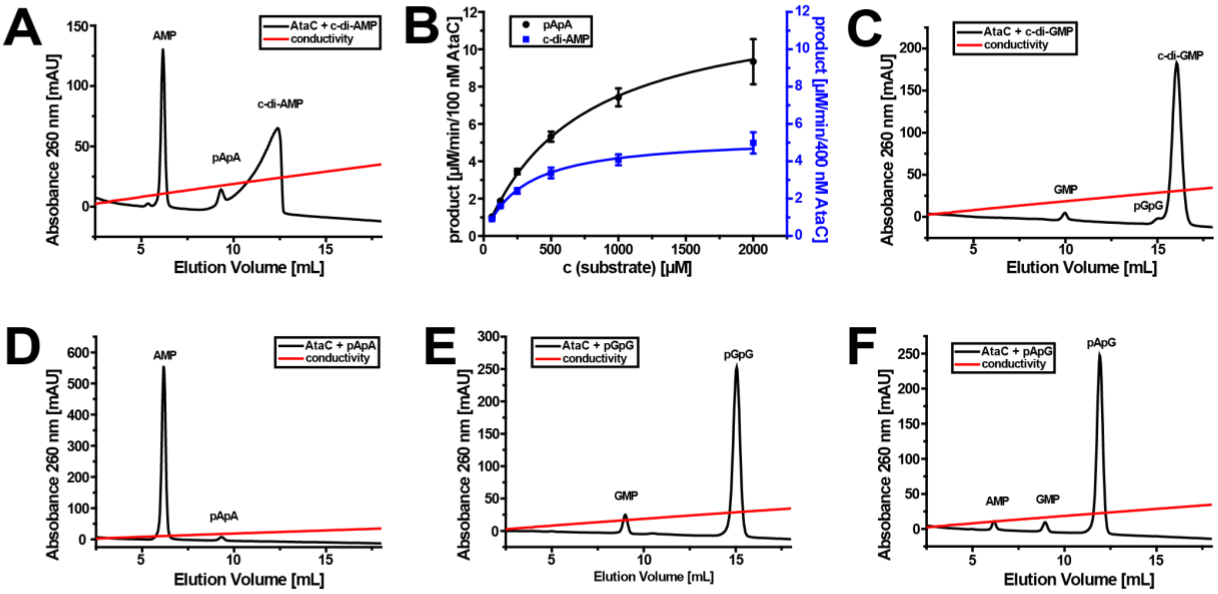


**Figure S1. Hydrolysis activity of AtaC is specific for adenosine bases.**

(A) Ion-exchange chromatography run on a Resource Q column of the reaction products after 1 h incubation from a 100 µl reaction containing 100 nM AtaC + 250 µM c-di-AMP.

(B) Michaelis-Menten kinetics of the reactions from 400 nM AtaC + c-di-AMP (62.5 – 2000 µM) and 100 nM AtaC + 5′-pApA (62.5 – 2000 µM) after 1 h of incubation at 37°C. c-di-AMP, $K_M$= 285 ± 32 µM, $k_{cat}$= 0.2 $s^{-1}$; 5′-pApA, $K_M$= 698 ± 32 µM, $k_{cat}$= 2.1 $s^{-1}$.

(C) Ion-exchange chromatography run on a Resource Q column of the reaction products after 1 h incubation from a 100 µl reaction containing 100 nM AtaC + 250 µM c-di-GMP.

(D) Ion-exchange chromatography run on a Resource Q column of the reaction products after 1 h incubation from a 100 µl reaction containing 100 nM AtaC + 250 µM c-di-AMP.

(E) Ion-exchange chromatography run on a Resource Q column of the reaction products after 1 h incubation from a 100 µl reaction containing 100 nM AtaC + 250 µM 5′-pGpG.

(F) Ion-exchange chromatography run on a Resource Q column of the reaction products after 1 h incubation from a 100 µl reaction containing 100 nM AtaC + 250 µM 5′-pApG.

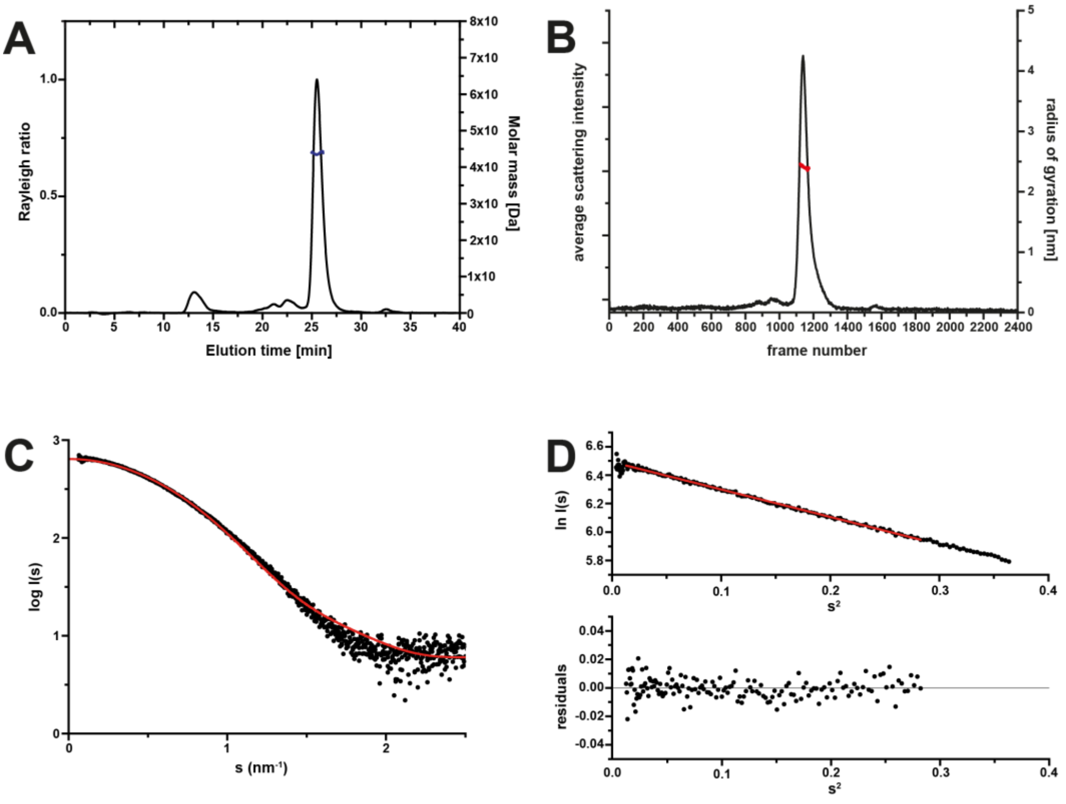


**Figure S2. AtaC is a monomer in solution.**

(A) Molecular weight determination of AtaC by SEC coupled multi-angle laser light scattering. The obtained molecular weight is 43.7 kDa and stable for the main protein peak at 25 ml.

(B) shows the relative scattering intensity of the sample during a size-exclusion coupled SAXS run at EMBL-P12 using a 24 ml Superdex increase S200 10/300 column (Intensity vs. frame No.). The respective estimated radius of gyration for each frame in the main peak is shown in red (right Y-axis).

(C) Measured SAXS curve of AtaC and a theoretical scattering curve (red) of the model of AtaC using PhnA as template (obtained from HHpred/MODELLER (21), $\chi_2$=3.6).

(D) Guinier plot ln I(s) vs. $s^2$ (top part) of the averaged buffer corrected scattering data (from B) and the respective residuals of the linear regression ($R_G$= 2.41 ± 0.1nm). The equally distributed errors of the linear regression (for s*$R_G$<1.3, Guinier approximation) indicates that the sample is not aggregating.

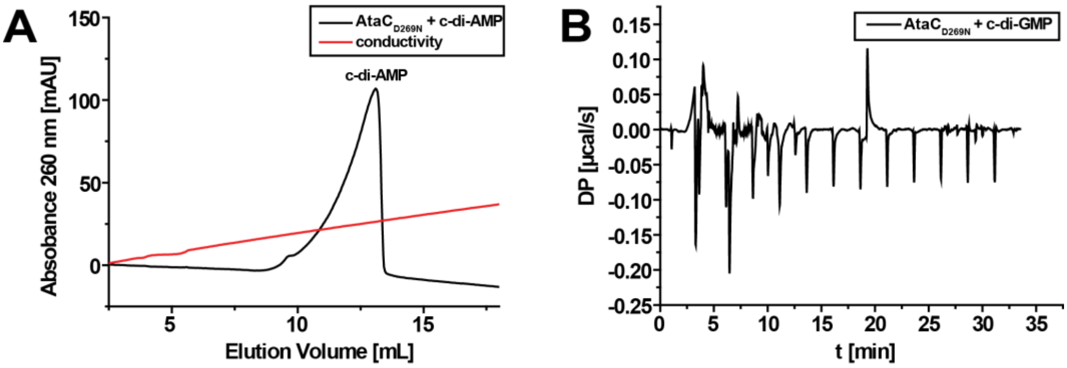


**Figure S3. AtaC$_{D269N}$ does not cleave c-di-AMP and does not bind c-di-GMP.**

(A) Ion-exchange chromatography run on a Resource Q column of the reaction products after 1 h incubation from a 100 µl reaction containing 1 µM AtaC$_{D269N}$ + 250 µM c-di-AMP.

(B) ITC measurement of 20 µM AtaC titrated with 140 µM c-di-GMP. No binding was detected.

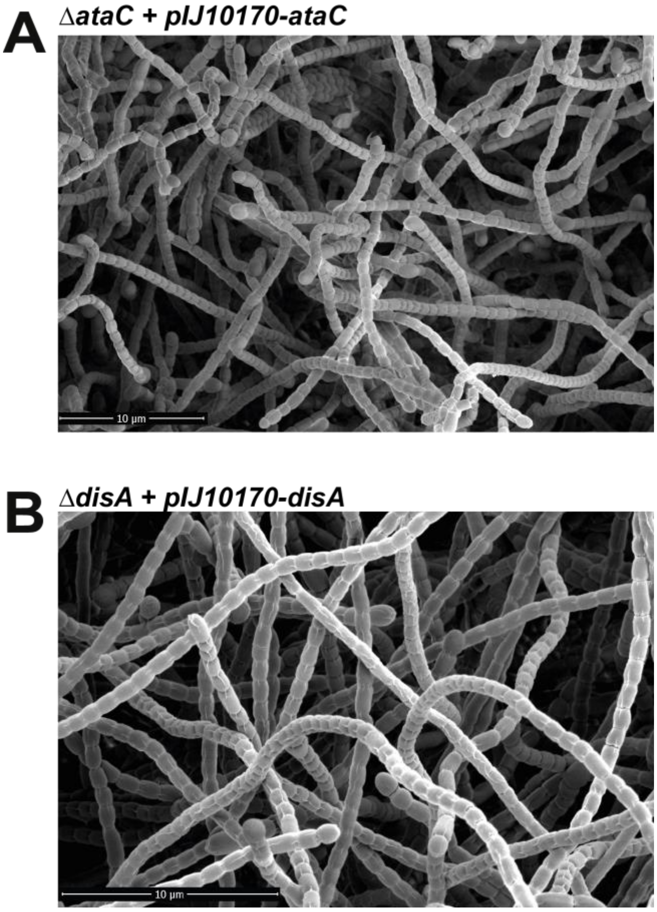


**Figure S4. Complementation of the Δ*ataC* and Δ*disA* mutants with the wild type alleles.** (A) Scanning electron micrographs show that expression of *ataC* from the *attB*$_{\Phi BT1}$ site under the control of the native promoter from pIJ10170 complemented the delayed developmental phenotype of the Δ*ataC* mutant. (B) Expression of *disA* from pIJ10170 in the Δ*disA* mutant did not alter the wild type phenotype of the mutant. For comparison see also Figure 5B in the main text. Cells were grown on MYM for 4 days at 30 °C.
